# Stochastic Forces in Microbial Community Assembly: Founding Community Size Governs Divergent Ecological Trajectories

**DOI:** 10.1101/2025.08.09.669462

**Authors:** Ibuki Hayashi, Martina Sánchez-Pinillos, Hirokazu Toju

## Abstract

Biological community dynamics are organized by the interplay of deterministic and stochastic processes. While species’ responses to biotic and abiotic environments determine the attractors of community compositions, stochastic processes, especially those in early stages of community assembly, can drastically influence the direction of temporal ecological dynamics. Nonetheless, establishing quantitative insights into the roles of stochastic processes has remained a major challenge. By developing a multi-replicated experimental system for tracking community assembly, we quantitatively evaluated the extent to which stochasticity could cause the divergence of community dynamics. We constructed soil- and freshwater-derived experimental microbial assemblages (microbiomes) with varying foundation community size, thereby controlling variation in initial community compositions among replicate communities (i.e., initial stochasticity). An analysis of > 3,000 community samples across four time points then indicated that higher levels of initial stochasticity could result in greater divergence of population- and community-level consequences. From the quantitative perspective, conspicuous differentiation into alternative trajectories of community assembly occurred when the absolute number of founding prokaryotic cells was less than the order of 10^4^. Furthermore, the replicate microbiomes differed in dominant *Pseudomonas* species, suggesting divergence into alternative transient/stable states through competitive exclusion between species occupying similar niches. Overall, our experiment showed that quantitative differences in stochasticity could result in qualitative differences in the fate of ecological communities. Further quantitative insights into stochasticity in community assembly will reorganize our views on biological invasions, agroecosystem microbiome management, and therapeutics of human-associated microbiomes.

## INTRODUCTION

The processes driving community assembly are broadly categorized into deterministic and stochastic ones (Clark 2009; Chase & Myers 2011; Vellend *et al*. 2014; Zhou & Ning 2017; Shoemaker *et al*. 2020). The structure of a community is primarily shaped by “selection” processes such as environmental filtering (Kraft *et al*. 2015) and interactions between species (HilleRisLambers *et al*. 2012), both of which impose deterministic impacts on community ecological dynamics. On a “ball-in-cup” analogy of community assembly, the deterministic (selection) processes are represented by the architecture of a landscape depicting the relationship between community-scale structure and stability (Fig. 1A right) (Beisner *et al*. 2003; Ridolfi *et al*. 2007; Van Meerbeek *et al*. 2021). Along with the selection processes, stochastic fluctuations in community compositions operate in community dynamics (Arani *et al*. 2021; Hastings *et al*. 2021). For example, ecological drift, which results from random demographic events occurring irrespective of species’ traits or niches, adds complexities to temporal changes in community structure (Hubbell 2005; Vellend *et al*. 2014; Shoemaker *et al*. 2020). On a stability landscape, such stochastic processes are illustrated as the fluctuations of community states, which may eventually diverge into different basins representing alternative stable states (Fukami 2015; Abbott & Nolting 2017; Hayashi *et al*. 2024) of community compositions (Fig. 1A left) (Beisner *et al*. 2003). Thus, fundamental insights into community assembly are available only when we understand the interplay between deterministic processes (changes in landscape architecture) and stochastic processes (random demographic fluctuations) (Jeraldo *et al*. 2012; Vellend *et al*. 2014; Måren *et al*. 2018; Ning *et al*. 2020).

**FIGURE 1.**
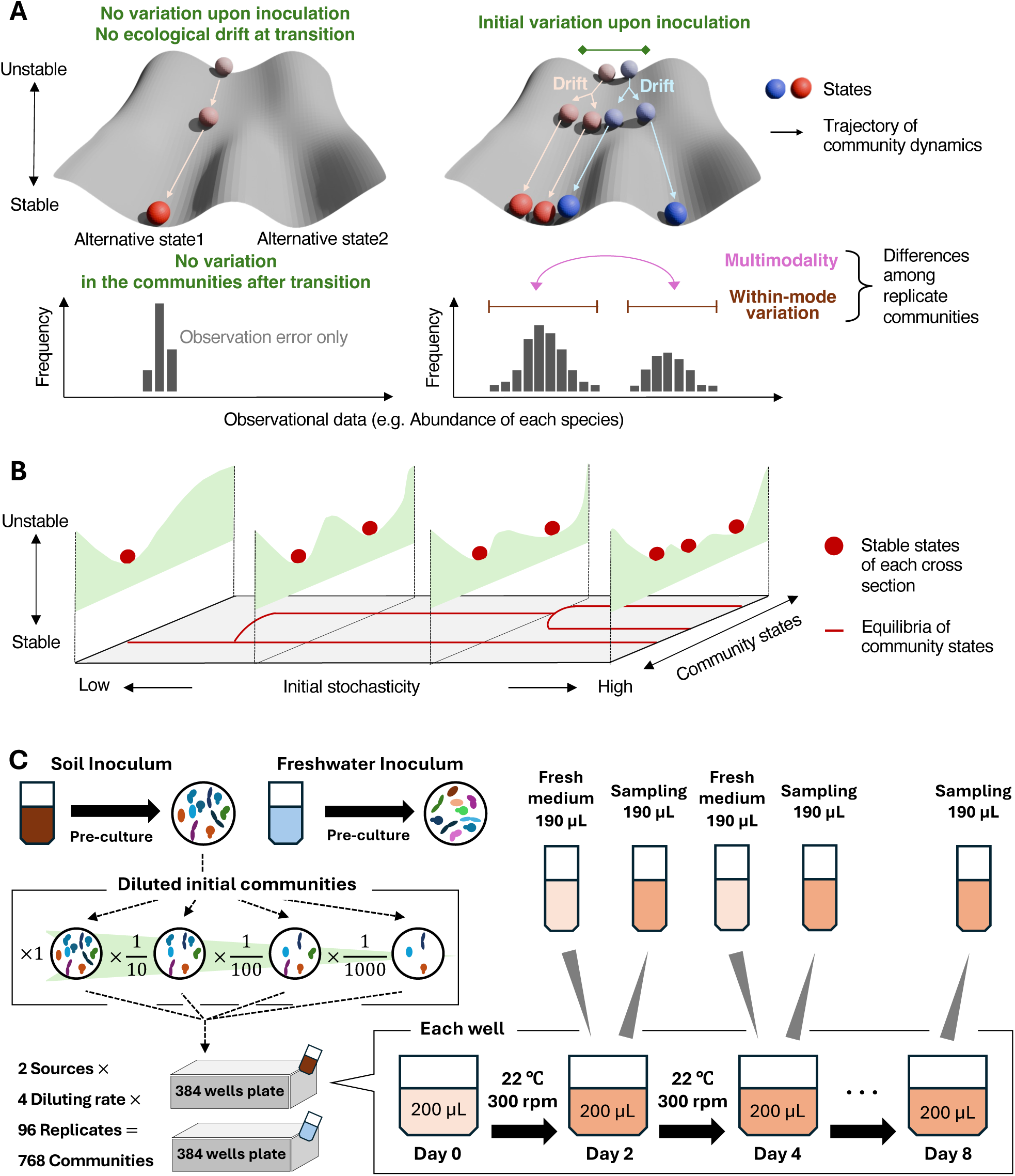
Theoretical concepts and experimental design. (A) Interplay of deterministic and stochastic processes (Framework 1). In fully deterministic ecological processes, divergence into alternative community states do not occur (left panel). In real ecosystems, divergence into alternative “basins” of attraction would occur on a stability landscape depending on stochasticity at the onset of a community (initial stochasticity) and ecological drift during community assembly. (right panel). Among-replicate variation in community structure can be quantified based on a histogram as detailed in Figures 3 and 4. (B) Initial stochasticity as a hyperparameter (Framework 2). In this alternative conceptual framework, initial stochasticity organizes the architecture of the stability landscape. For example, bifurcations (i.e., changes in the number of attractors) may occur along an axis of initial community density. (C) Experimental design. Each microbiome sample derived from a soil or freshwater ecosystem was precultured with cycloheximide at room temperature for 2 days to remove eukaryotes. Each inoculum source was diluted into four dilution series, yielding eight inoculum settings (2 sources × 4 dilution series). For each inoculum setting, 96 replicate experimental communities were constructed. A fraction of the culture fluid was sampled every 2 days (48 hours), and an equivalent volume of fresh medium was added to the continual culture system throughout the eight-day experiment (8 inoculum settings × 96 replicates × 4 time points = 3,072 community samples).

While possible destinations of community dynamics (system’s attractors) *per se* are set by deterministic processes, stochastic processes can determine the direction of community assembly (i.e., divergence into alternative basins). In particular, at early, unstable phases of community assembly, slight differences in community compositions can have great impacts on the subsequent community dynamics, causing divergence into alternative basins of attraction (Fig. 1A right) (Zhou *et al*. 2013; Abbott & Nolting 2017; Lerch *et al*. 2023; Hayashi *et al*. 2024). Without stochasticity during the initial stages of community assembly, repeated observations under identical external conditions would be expected to yield identical communities (i.e., completely deterministic assembly; Fig. 1A, left). Importantly, the degree to which communities diverge into alternative states may depend on the level of initial stochasticity (Scheffer & Carpenter 2003; Lopes *et al*. 2024). The community size of founding species assemblages, for example, would limit the range of the subsequent fluctuations in community dynamics, thereby controlling bifurcations into alternative basins (Abbott & Nolting 2017). Nonetheless, few empirical studies have examined how stochastic components in early community assembly generate varied consequences. We still have limited knowledge of how the “multi-stable” patterns of ecological communities could be realized depending on the level of initial stochasticity.

In comprehensively understanding community dynamics on stability landscapes, it is essential to obtain quantitative experimental insights into stochastic processes (Shoemaker *et al*. 2020). In principle, the quantification of stochasticity requires repeated observations. Therefore, if we are to evaluate the effects of stochastic processes in community assembly, it is crucial to obtain bird’s-eye views on the fates of many replicated communities (Pascual-García *et al*. 2025). As discussed above, the existence of multiple basins does not guarantee that alternative community states will actually be observed (Spagnolo *et al*. 2003; Mankin *et al*. 2006). Thus, it is crucial to examine how the level of divergence into alternative basins depends on initial stochasticity in community assembly. In experimental systems with many replicates, we can control stochasticity by changing the size of founding communities, while keeping the contributions of deterministic (selection) processes (Siqueira *et al*. 2020) (Fig. 1A right). In other words, decreased initial community size is expected to increase among-replicate variability in early assembly (i.e., initial stochasticity), resulting in varied consequences in later stages. In systematically conducting such experiments, microbial communities (microbiomes) serve as ideal research targets because they allow us to design experiments with tens of replicated communities, among which we can evaluate varied consequences (e.g., variation in destination basins) (Estrela *et al*. 2022; Hayashi *et al*. 2024).

Moreover, since generation cycles are much shorter in microbes than in macro-organisms (e.g., plants and animals), community experiments with microbes can be performed in short time periods. Consequently, highly replicated microbiome experiments with varying initial community sizes will provide quantitative insights into the diversification of community dynamics.

In this study, we constructed a highly replicated experimental system to examine how initial community size determines the divergence into alternative community states. We established laboratory microbiomes using two types of source communities (soil and freshwater source communities), each of which was subjected to a dilution series with four levels. For each dilution level, 96 replicate communities were tracked across four time points. The community compositional analysis of the > 3,000 samples (2 source community types × 4 initial concentration levels × 96 replicates × 4 time points) allowed us to examine how initial stochasticity level could drive community assembly. We hypothesized that lowering inoculum concentrations would increase the level of initial stochasticity, which, in turn, could amplify variation in community composition during subsequent assembly processes. The hypothesis was tested by calculating two types of indices that allowed the quantification of among-sample divergence of community structure.

Specifically, across the 96 replicates at each time point, we evaluated multi-peak patterns in the frequency distribution of each species’ abundance as indicators of divergence into alternative basins of attraction. We also quantified the variation within each peak (“within-mode variation”) as a measure of fluctuations within each basin (Fig. 1A, right). We then tested the relationship between initial stochasticity level and each index representing the diversity of realized ecological patterns.

Furthermore, by applying the “ecological dynamic regime” framework for time-series community data (Sánchez-Pinillos *et al*. 2023), we quantitatively examined how founding community size could determine the diversity of the trajectories of community dynamics. Overall, the systematic experimental design with a high number of replicate communities provides quantitative insights into the mechanisms underlying the formation of alternative communities.

## MATERIALS AND METHODS

### Concepts and assumptions

In considering how initial stochasticity level influences community stability, there can be alternative conceptual frameworks for interpreting observed patterns. One is to assume a fixed stability landscape structure on which the community dynamics are driven by stochastic processes (Framework 1). In this assumption, the landscape structure remains unchanged with initial stochasticity, and temporal dynamics vary among replicate communities depending on starting positions (compositions at the onset of communities) and the subsequent fluctuations on the fixed landscape (Fig. 1A right). The other way of interpreting community assembly data is to assume variation in stability landscape topography across different initial stochasticity levels (Framework 2). In this alternative assumption, initial stochasticity is treated as a “hyperparameter” controlling the architecture of stability landscapes, which is represented by the number of attractors (Fig. 1B).

In this study, we interpret community assembly data mainly based on Framework 1. We quantify among-replicate variation in community compositions based on a statistical distribution model. We then examine whether the initial stochasticity level influences variation in population- and community-level consequences. Meanwhile, to interpret the data based on Framework 2, we apply recently proposed statistical frameworks assuming the dynamic nature of stability landscapes (Suzuki *et al*. 2021; Masuda *et al*. 2025).

### Continuous culture of microbiomes

While “synthetic community” approaches using explicitly defined sets of microbial species or strains have been commonly applied, experiments with field-collected assemblages of diverse taxa are expected to provide more realistic insights into prokaryotic community assembly. We collected two types of field-collected microbiomes as source communities of the experiment. One is sampled from the soil of the A layer (0-10 cm in depth) in the research forest of Center for Ecological Research, Kyoto University, Shiga, Japan (34.972 °N; 135.958 °E) on January 30, 2023. The other source microbiome derived from the surface water of a freshwater pond near the Center for Ecological Research (34.974 °N, 135.967 °E). After sampling, the soil was sieved with a 4-mm stainless mesh and then 5 g of the sieved soil was mixed in 40 mL sterilized PBS buffer (the detail is shown in Table S1). The freshwater sample was sieved with 20 μm filter and then 20 mL of the filtered water was mixed in 20 mL sterilized 2 × PBS buffer. In the preparation procedures, we added cycloheximide at a concentration of 200 μg/mL to exclude eukaryotes from the source microbiome. The source prokaryote microbiome was cultured at 22 °C for 48 hours and diluted 10 times to make inoculum solutions for the subsequent experiment.

We then quantified the number of prokaryotic cells in the source communities (inoculum prokaryotic cell suspensions) based on a quantitative DNA sequencing platform as detailed below (see the subsection “DNA metabarcoding”). To make a gradient of initial community size, the base inoculum sample (×1) was serially diluted for three steps, making ×1/10, ×1/100, and ×1/1000 inocula for each of the soil and freshwater sources (Fig. 1C). We introduced each of the eight inoculum microbiomes (2 sources × 4 dilution rates) into a complete artificial medium with 96 replicates (in total, 8 inoculum settings × 96 replicates = 768 experimental communities). To make the compositions of the media as simple as possible, we used M9 medium with minimal inorganic additives and three types of carbon resources (glucose, leucine, and citrate as detailed in Table S1). In each well of a 240 μL deep-well plate, 10 μL of the inoculum microbiome solution and 190 μL of medium were installed. The 384 deep-well plate was kept shaken at 200 rpm using a plate thermo-shaker BSR-MB100-4A (Bio Medical Sciences Co. Ltd., Tokyo) at 30 °C for two days. After two-days incubation, 190 μL out of the 200-μL culture medium was sampled from each of the 96 wells after mixing (pipetting) every two days for 8 days. All pipetting manipulations were performed with high precision using a 384-channel automatic pipetting machine (EDR-384SR, BIOTEC Co. Ltd., Tokyo) placed in a sterilized environment within a laminar flow cabinet. In each sampling event, 190 μL of fresh medium was added to each well so that the total culture volume was kept constant. In total, 3,072 samples (768 communities/day × 4 time points) were collected.

### DNA metabarcoding

The community compositions of the microbiome samples were analyzed based on the DNA metabarcoding of the 16S ribosomal RNA (rRNA) region based on a previously reported protocol of DNA extraction, PCR, purification, library preparation, and Illumina sequencing (Hayashi *et al*. 2024) as detailed in Supplementary Methods.

The 16S rRNA sequencing analysis was applied to the source communities (soil and freshwater inoculum microbiomes) as well. In the DNA metabarcoding of the source communities, we used a quantitative amplicon sequencing platform for estimating the copy numbers of 16S rRNA in the template DNA aliquots (Ushio *et al*. 2018; Fujita *et al*. 2023a) (see “the PCR and DNA sequencing” section in Supplementary Methods for details).

### Bioinformatics

In total, 66,302,381 sequencing reads were obtained with the Illumina sequencing. The raw sequencing data obtained in the Illumina sequencing were converted into FASTQ files using the program bcl2fastq 1.8.4 distributed by Illumina. The output FASTQ files were then demultiplexed with the program Claident v0.9. 2022.01.26. The sequencing reads were subsequently processed with the program DADA2 (Callahan *et al*. 2016) v.1.18.0 of R 4.2.2 to remove low-quality data. The molecular identification of the obtained amplicon sequence variants (ASVs) was performed based on the naive Bayesian classifier method (Wang *et al*. 2007) with the SILVA v.138.1 database (Quast *et al*. 2013).

In the analysis of the experimental culture samples, the diversity of microbial ASVs saturated at the sequencing depth of 3,000 reads per sample (Fig. S1). Therefore, the sequencing data of the culture samples was rarefied to 3,000 reads per sample with the “rrarefy” function of the R vegan 2.6.6.1 package (Oksanen *et al*. 2025). Of the 3,072 samples (8 inoculum settings × 96 replicates × 4 time points), 3,063 samples with more than 3,000 reads were used in the following pipeline. After screening for the replicate communities for which sequencing data were available for all the four time points, 3,040 samples were subjected to the following statistical analyses. In total, 490 prokaryote ASVs belonging to two kingdoms, 26 classes, 64 orders, 88 families, and 147 genera were detected (Fig. 2). The ASVs were re-clustered into operational taxonomic units (OTUs) with 99% thresholds using the program VSEARCH v2.15.2 (Rognes *et al*. 2016), yielding 337 OTUs.

**FIGURE 2.**
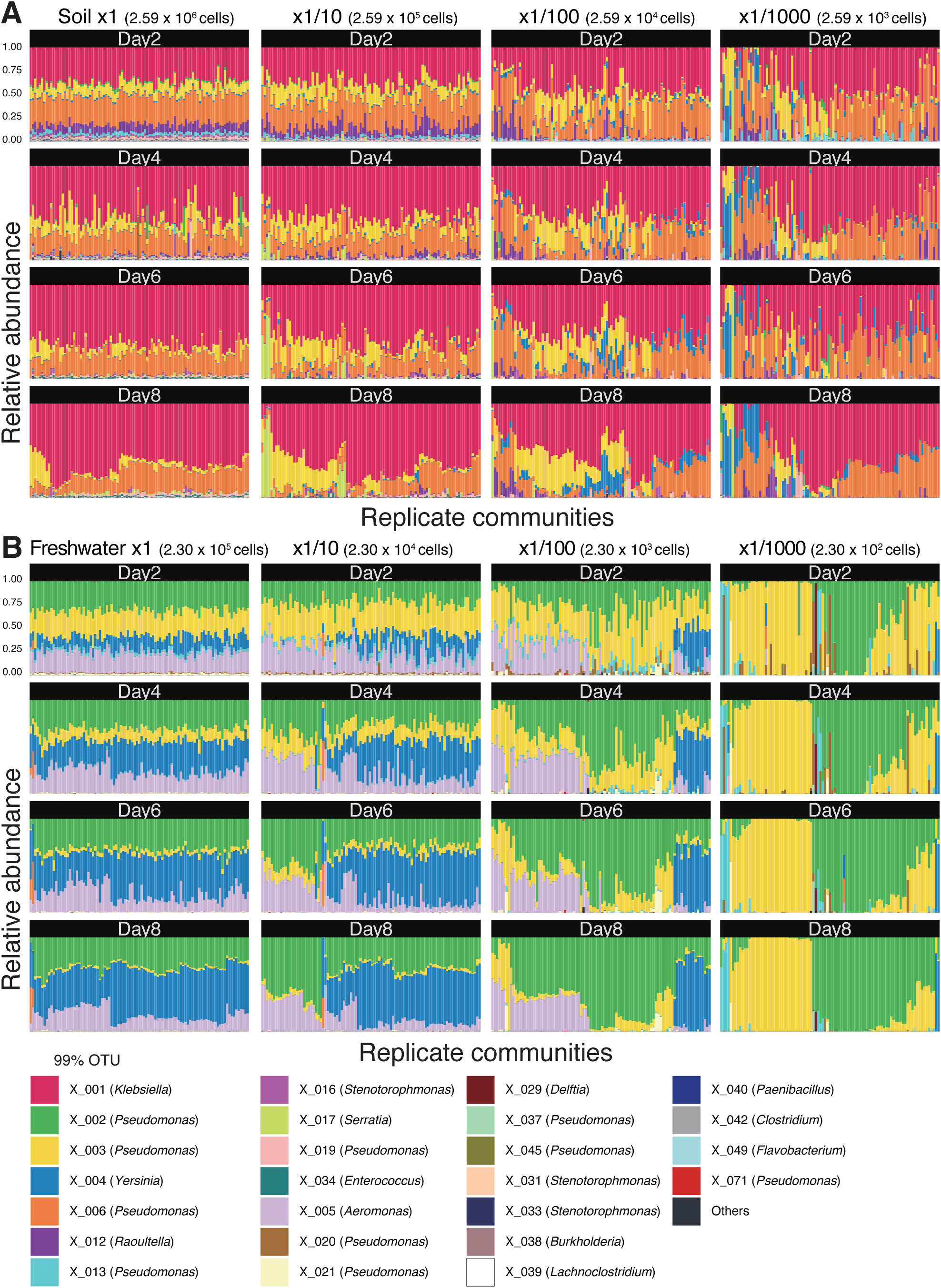
Overview of among-replicate variation in community structure. (A) Experiment with soil inoculum microbiome. Temporal changes in the 99% OTU-level community compositions (relative abundance) are shown. For each inoculum dilution rate, replicate communities were ordered based on the results of unweighted pair group method with arithmetic mean (UPGMA) analyses performed on Day 8. (B) Experiment with freshwater inoculum microbiome.

For the quantitative sequencing of the source communities, the estimated densities of 16S rRNA gene copies were obtained. The 16S rRNA gene copy estimates were then converted into the estimated number of prokaryotic cells introduced into the experimental microbiomes. In this calculation, the number of 16S rRNA tandem repeats within a genome was inferred for each taxon constituting the source communities based on the RasperGade16S database v0.0.1.0 (Gao & Wu 2023).

### Estimating initial variation upon inoculation

The data of each OTU’s cell numbers in the inoculum communities were used to quantify the level of stochasticity at the initial stage of the experimental microbiome assembly. Specifically, the cell number of each OTU introduced into each sample was assumed to follow a multinomial distribution, with parameters being the total number of cells on average in the inoculated cell suspension and the relative abundance of each taxon in the suspension. By assuming random picking of cells from the inoculum suspension, a simulation following the multinomial distribution was performed 1,000 times for each inoculation setting. We then estimated variance in introduced cell numbers among replicated culture samples for each prokaryotic OTU in each of the eight inoculum settings (2 sources × 4 dilution rates). For each OTU in each inoculum setting, the variance was used to calculate coefficient of variation (CV), which represents the level of initial stochasticity in the microbiome experiment (hereafter, initial variation upon inoculation; Fig. 3A).

**FIGURE 3.**
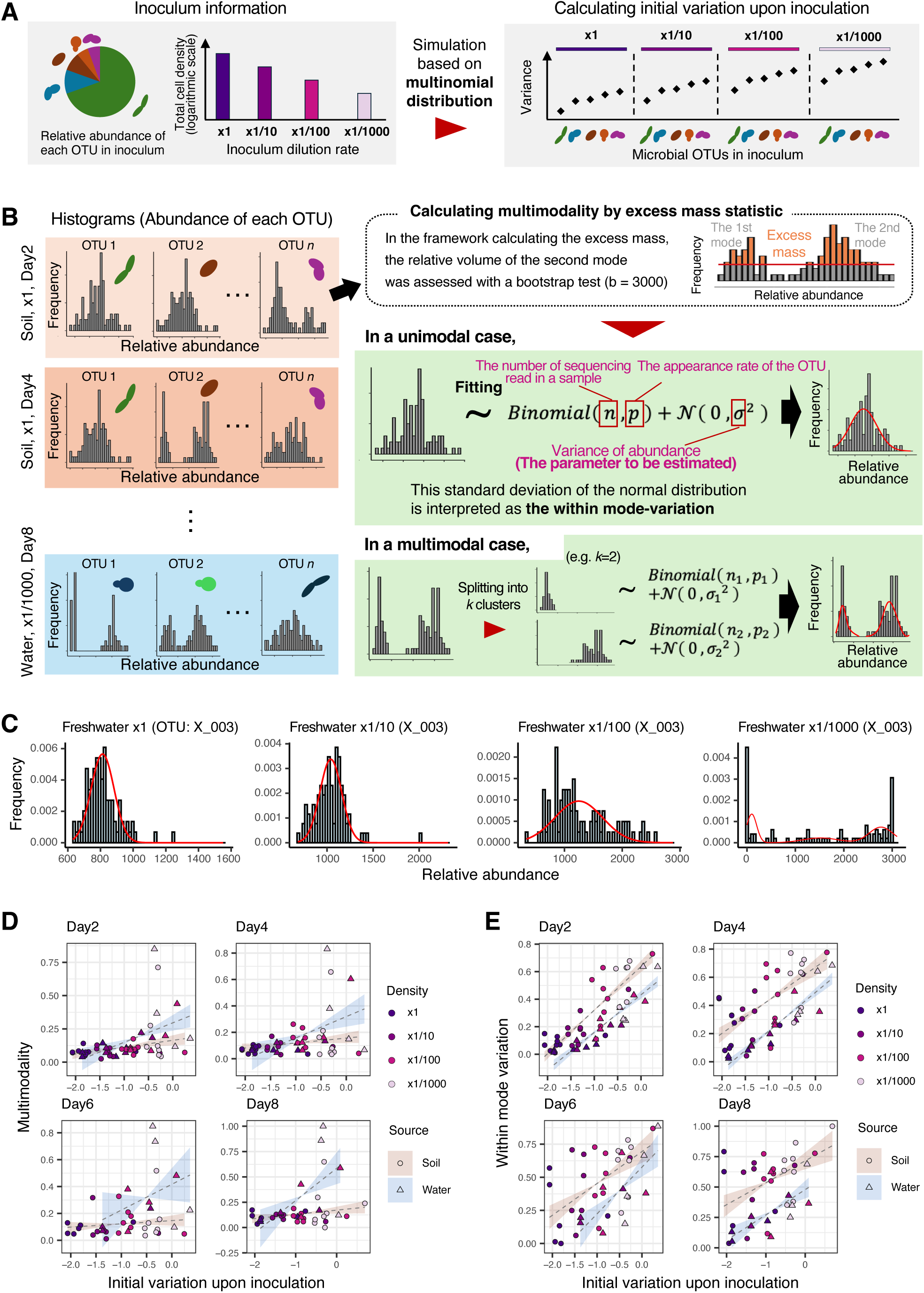
Initial stochasticity and divergence in OTU abundance among replicate communities. (A) Calculation of initial variation upon inoculation. When the relative abundance of each OTU and the total number of cells in the inoculum are known, the number of cells introduced for each OTU at inoculation follows a multinomial distribution, enabling calculation of the coefficient of variation (CV). The CV generally decreases with higher relative abundance and larger total numbers of inoculated cells. (B) Multimodality and within-mode variation of OTU abundance. For each microbial OTU, the degree to which its abundance histogram exhibits multiple peaks is assessed as multimodality. For each peak identified in the multimodality analysis, among-replicate variation was assumed to follow binomial and standard normal distributions. Within-mode variation is then evaluated as the standard deviation of the standard normal distribution. (C) Examples of abundance histograms. For an OTU (“X_003”) in the freshwater microbiome experiment, the histograms of read counts across replicate communities are overlaid with predictions from the mixture model (panel B). The data of Day 2 at each inoculum dilution rate are shown (see Fig. S6 for results on other OTUs). (D) Relationship between initial stochasticity and multimodality. By targeting the OTUs commonly observed across experimental replicates, relationship between initial variation upon inoculation (panel A) and multimodality was examined for each source microbiome type (soil or freshwater) for each time point. The multimodality estimates are scaled from 0 to 1 across the panels. (E) Relationship between initial stochasticity and within-mode variation. The within-mode variation scores are scaled from 0 to 1 across the panels.

### Multimodality in OTU abundance

For each combination of source microbiomes and inoculum dilution rates (inoculum setting), we examined the degree to which the abundance of each prokaryotic OTU varied among replicate community samples at each time point after the inoculation event. Based on the stability landscape concept of community assembly, the among-sample variation in OTU abundance was quantified sequentially with two types of indices (Fig. 1A right). For the first step, the presence of multiple peaks in the histogram of each OTU’s abundance across the replicate samples was examined with a “multimodality” index, which give insights into the presence of multiple basins of attraction in the microbiome assembly (Fig. 3B). Next, for each peak identified in the histogram, “within-mode” variation among replicate samples was calculated by assuming fluctuations within each basin of a stability landscape (Fig. 3B). We then examined whether each type of among-replicate variation could increase with increasing initial variation upon inoculation (i.e., stochasticity at the foundation of the replicate communities).

In the calculation of the multimodality, OTUs that appeared in more than ten communities and accounted for at least 0.5% of the total sequencing reads were targeted. In total, a total of 174 OTU frequency data collected across all time points were analyzed. A multimodality test (Ameijeiras-Alonso *et al*. 2019) was then performed to calculate excess mass statistic and *p-*values for each OTU in each inoculum setting at each time point. This scaled excess mass statistic obtained in the multimodality test was defined as the multimodality of each OTU’s abundance. The multimodality test was conducted with the R package multimode 1.5 (Ameijeiras-Alonso *et al*. 2019) by setting the number of replicates to 3,000. The *p-*values indicating the presence/absence of multiple peaks (modes) were adjusted for multiple comparisons at each time point based on false discovery rate (FDR) to obtain *q*-values.

### Within-mode variation in OTU abundance

We next calculated “within-mode” variation, which is expected to reflect the level of fluctuation within each basin of attraction in the post-inoculation processes (Fig. 1A right). To calculate the additional measure of among-sample variation, we identified the number of peaks in the histograms of OTU abundance. Specifically, we classified OTUs with unimodal and multimodal distributions by applying the abovementioned multimodality test with a threshold *q*-value (significance level) of 0.05 in each inoculum setting at each time point. The frequency of OTU abundance within each peak (mode) was then assumed to follow a mixture model of a binomial distribution and a standard normal distribution (Fig 3B). In principle, by the parameters of the binomial distribution were automatically obtained as the number of trials (i.e., the rarefied number of sequencing reads per sample = 3,000) and the probability of observations (i.e., the relative abundance of a target OTU in a target community sample) based on the assumption of Bernoulli trial. Therefore, the standard deviation within the normal distribution part in the mixture model was used as a measure of among-sample variation caused after the inoculation event.

The fitting to the mixture model was conducted by optimizing following likelihood function:

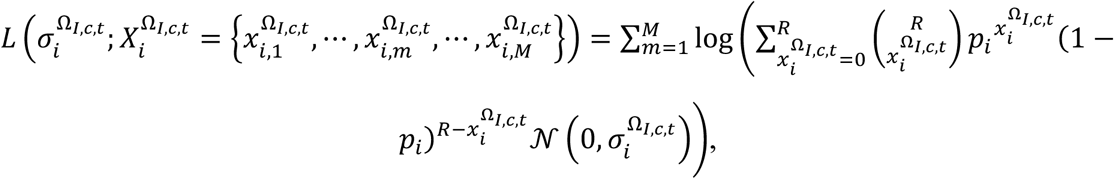

where 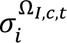 was the standard deviation of a standard normal distribution, 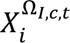 indicated the sequencing read numbers of OTU *i* in each subset data Ω [subscripts *I*, *c*, and *t* denoted the type of inoculum communities (soil or freshwater source communities), the dilution rate of inoculum community, and time point, respectively], and *M* was the number of replicate communities. In the right hand of the equation, *R* denoted the threshold number of sequencing reads (3,000 reads/sample in the rarefaction step of this study), *p_i_* was the relative abundance of OTU *i* of each subset Ω, and *N*(0, σ) indicated a standard normal distribution whose standard deviation was σ. The standard deviation 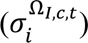 was estimated by minimizing the likelihood function using the “optimize” function in R. To gain reliable estimates, we conducted the model fitting ten times and then used the mean value of the estimates as “within-mode” variation for each peak of a target OTU in each inoculum setting at each time point. For each of the OTUs with multiple peaks, a mean value of the estimates was calculated. The within-mode variation estimate was then scaled by dividing it by the mean abundance of each OTU. Since the estimates of standard deviations were highly influenced by the presence of outliers, five of the largest and five of the smallest outliers were removed (see Table S2 for the information of the removed samples).

For the analysis of the OTUs with multimodal distributions, the data were divided into an optimal number of clusters. Specifically, after applying *k*-means clustering, the optimal number of clusters was inferred based on the silhouette coefficients calculated with the cluster 2.1.6 package (Maechler *et al*. 2024) of R. The clusters that contained six or more replicate samples were used in the fitting to the mixture model: outliers were not removed in the analysis for the multimodal cases.

### Community-scale differentiation among replicates

By extending the statistical approach applied at the OTU level analysis, we next developed a method for quantifying community-level differentiation of community structure. Instead of the distribution of each OTU’s abundance across samples (Fig. 3B), we focused on the distribution of pairwise dissimilarities (Bray-Curtis *β*-diversity) between replicate samples (Fig. 4A-B). The multimodality of pairwise community dissimilarity distributions was calculated for each inoculum setting at each time point to evaluate the extent to which replicate communities diverged into multiple basins of community structure.

**FIGURE 4.**
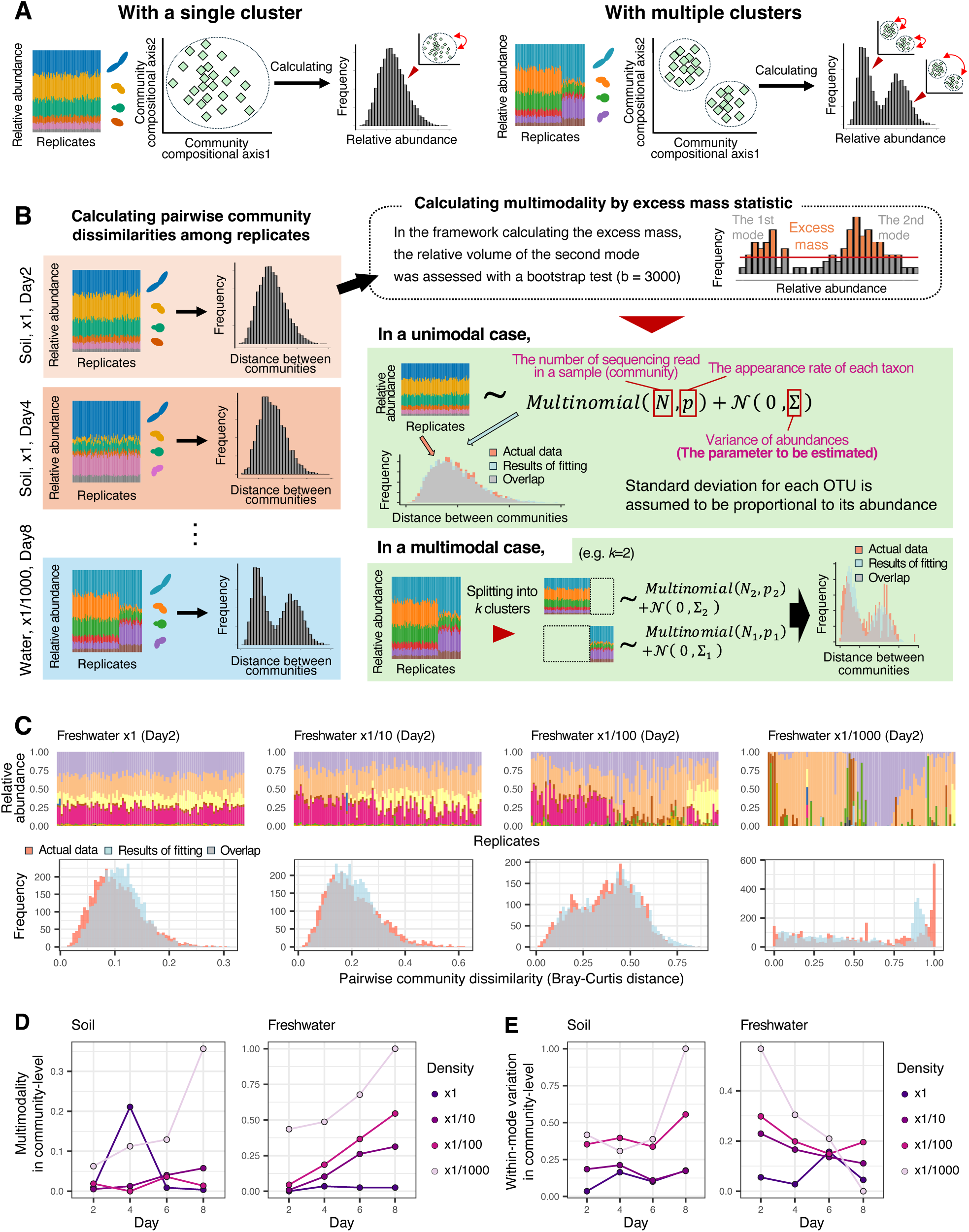
Among-replicate divergence in community compositions. (A) Patterns in pairwise community dissimilarity and underlying community structure. If replicate communities are divided into clusters according to their compositions, multiple peaks would appear in the histogram of among-replicate community dissimilarity. (B) Multimodality and within-mode variation in community dissimilarity histograms. The degree to which the histogram of pairwise community dissimilarity exhibits multiple peaks is assessed as community-scale measure of multimodality. For each peak identified in the multimodality analysis, among-replicate variation was assumed to follow multinomial and standard normal distributions. Within-mode variation is then evaluated as the standard deviation of the standard normal distribution. (C) Examples of community dissimilarity histograms. For freshwater-derived experimental microbiomes, the histograms of the histograms of community dissimilarity among replicates are overlaid with predictions from the mixture model (panel B). The data of Day 2 at each inoculum dilution rate are shown (see Fig. S8 for full results). (D) Temporal changes in multimodality. For each combination of source microbiome type (soil or freshwater) and inoculum dilution rate, the temporal trends of community-scale multimodality are shown. Multimodality estimates are scaled from 0 to 1 across the panels. (E) Temporal changes in within-mode variation. The community-scale estimates of within-mode variation are scaled from 0 to 1 across the panels.

Likewise, variation in community structure within each basin of the stability landscape (Fig. 1A) was inferred by quantifying within-mode variation in the distribution of pairwise community dissimilarity. For each inoculum setting at each time point, we checked whether multiple peaks existed within the histogram of pairwise community dissimilarity. In general, the presence of multiple peaks within the distribution of pairwise dissimilarities indicates that the focal community dataset includes multiple clusters (groups) of data points (Hayashi *et al*. 2024). In our analysis, if the presence of multiple groups was supported in a multimodality test (threshold *q*-value = 0.05; Fig. 4B), we performed a *k*-means clustering analysis followed by a silhouette coefficient analysis to split the replicate communities into groups.

For each split dataset, the community compositions of replicate samples were assumed to follow the following mixture model of a multinomial distribution and a standard normal distribution:

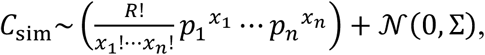

where *C_sim_*. was the community matrix derived from the mixture model. In the multinomial part 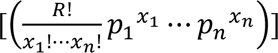, *R* was the threshold number of sequencing reads (= 3,000 reads/sample), *x_i_* is the random variable of multinomial distribution corresponding to the read count of OTU *i* 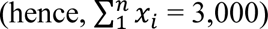, and *p_i_* was a parameter of multinomial distribution corresponding to the occurrence probability of OTU *i*. In the part of normal distribution [*N*(0, Σ)], standard deviation of each OTU was assumed to be proportional to its occurrence probability (*p_i_*) as follows:

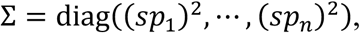

where *s* is a constant representing the variability of community structure among replicate samples. In the simulation with the mixture model, the occurrence probability of each OTU (*p_i_*) was estimated from the abundance information of each OTU within the original data matrix by assuming a Dirichlet distribution with the “rdirichlet” function in the R package MCMCpack 1.7.1 (Martin *et al*. 2024). The most likely variability constant (*s*) in the normal distribution part were determined by minimizing the difference between the distribution derived from the actual data and the distribution derived from the mixture model as follows:

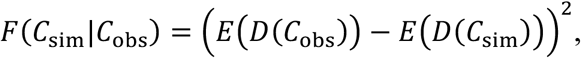

where *C*_sim_ was the community matrix generated by the mixture model and *C*_obs_ was the community matrix of the observed data. In the right hand of the equation, *E*(*D*(*C*_obs_)) represented an empirical cumulative distribution function (eCDF) of Bray-Curtis dissimilarity among replicate samples in the observed data matrix *C*_obs_, while *E*(*D*(*C*_sim_)) was an eCDF of Bray-Curtis dissimilarity in the simulated data matrix *C*_sim_. This solving process was conducted one hundred times, among which the run with the smallest *F*(*C*_sim_|*C*_obs_) was used for gaining a reliable estimate of the constant (*s*) representing community variability within a basin of community structure.

### Ecological dynamic regimes

In the analysis of multimodality and within-mode variation, we assumed the presence of a fixed stability landscape for each of the soil-derived and freshwater-derived experimental communities (Framework 1: Fig. 1A). An alternative approach to interpret the community data is to assume that initial stochasticity determines the topography of stability landscapes *per se* (Framework 2: Fig. 1B). To visualize how stability landscape architecture changes with initial stochasticity level, we performed a statistical-physics-based approach for inferring the relationship between systems’ states and their probabilities (Suzuki *et al*. 2021; Masuda *et al*. 2025). For each of the eight inoculum settings, we applied the energy landscape analysis (Suzuki *et al*. 2021) for statistically modeling the probabilities of observing given community compositions (Fujita *et al*. 2023a, 2025; see Supplementary Information for the details of the statistical approach).

We further evaluated whether the stochasticity level at the initial stage of community assembly could determine changes in community dynamic regimes. To quantify how the trajectories of community assembly diverged over time among replicate samples, we applied the “ecological dynamic regime” (EDR) framework for time-series community data (Sánchez-Pinillos *et al*. 2023). This statistical framework has been developed for identifying the dynamic property of steady states, which are represented by divergence into alternative transient/stable states and fluctuations within basins of attraction. Based on the time-series data of community compositions, the EDR framework allows characterizing and comparing ecological dynamic regimes in a multidimensional state space defined by the temporal changes in replicate communities represented by trajectories. Thus, we defined a dynamic regime as the group of trajectories (i.e., time series) corresponding to each source (soil or freshwater) and dilution rate. As detailed in Sánchez-Pinillos *et al*. 2023, a “representative trajectory” passing by the densest regions of the state space (i.e., containing many trajectories) can be used to summarize the main dynamic patterns of observed community assembly. The statistical approach then provides three metrics for quantifying the distribution of ecological trajectories among replicate communities defining a dynamic regime.

The dynamic dispersion (dDis) quantifies the average dissimilarity of each trajectory from the representative trajectory. The dynamic dispersion index is calculated as follows:

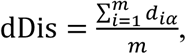

where *d_iα_* represents dissimilarity between a trajectory *i* and a reference trajectory *α*, and *m* is the number of observed trajectories (i.e., the number of replicate time series). When quantifying the dispersion of community dynamics from the overall direction of assembly, we selected the representative trajectory with the highest resolution of state space, based on the average depth of representative trajectories, as the reference trajectory. Alternatively, a reference trajectory could be selected based on other criteria, such as the one with the largest number of segments (selection by size), the one with the smallest average link dissimilarity connecting segments within the trajectory (selection by average link), or the one whose constituent segments best represent all segments in the EDR on average (selection by average density). To evaluate these alternatives, we conducted calculations using (i) the representative trajectory that performed best according to each of the size, average link dissimilarities, and average density metrics, and (ii) the trajectory with the lowest total rank across all four metrics: average depth, size, average link, and average density. The overall trends did not differ substantially from those obtained using average depth as the selection criterion. The dynamic dispersion score varies from 0 (when all trajectories are identical to the representative trajectory) to 1 (when replicate communities take completely different trajectories of community dynamics).

The dynamic beta diversity (dBD) quantifies the average dissimilarity between community trajectories as follows:

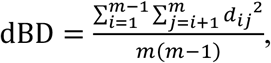

where *d_i,j_* denotes dissimilarity between trajectories *i* and *j*. Like the dynamic dispersion, the dynamic beta diversity varies from 0 (completely identical dynamic among replicates) and 1 (complete differentiation of temporal community dynamics).

The dynamic evenness (dEve) measures the continuity of trajectory variation within the dynamic regime as follows:

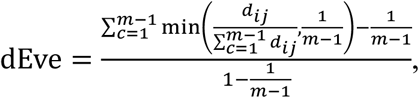

where *c* is the edges of a minimum spanning tree constructed from the set of trajectories forming a dynamic regime. In contrast to the dynamic dispersion and dynamic beta diversity, a higher value of dynamic evenness indicates a lower level of divergence among the time series of replicate communities. It ranges from 0 (when many subclusters of trajectories exist) and 1 (when all trajectories are evenly distributed).

The three EDR metrics were calculated for each of the eight inoculum settings (2 source communities × 4 dilution rates) defining the dynamic regimes using the R ecoregime 0.2.0 package (Sánchez-Pinillos *et al*. 2023). In addition, the representative trajectories of temporal community dynamics were inferred with the ecoregime package (the minSegs parameter in the ‘retra_edr’ function was set as 5). For each inoculum setting, the representative trajectories are shown on a two-dimensional surface of community compositions, which was defined based on a principal coordinate analysis (PCoA) using Bray-Curtis dissimilarities (Fig. 5).

**FIGURE 5.**
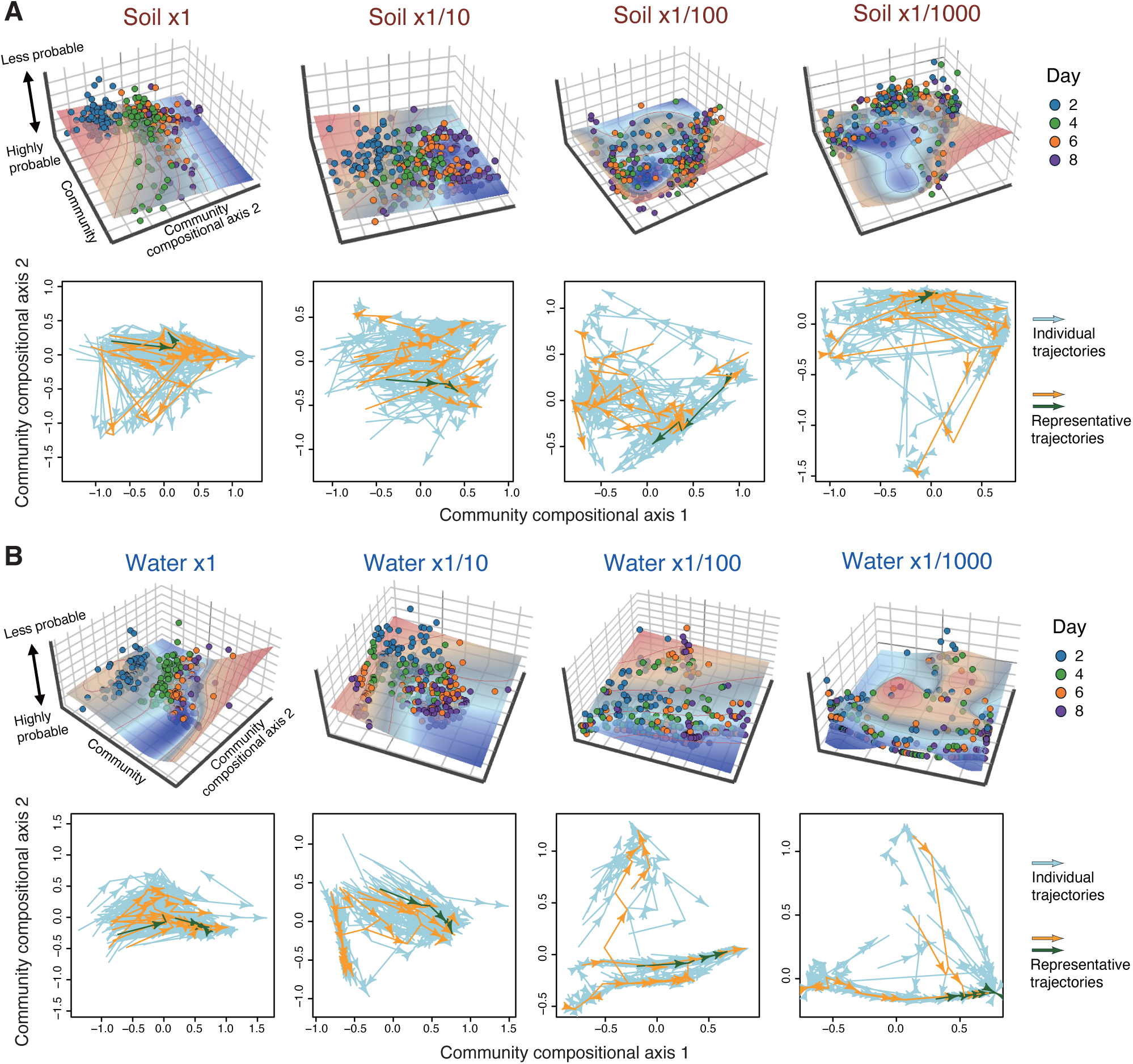
Trajectories of community dynamics. For each combination of source microbiome type (soil or freshwater) and dilution rate, the energy landscape analysis was performed to infer the topography of a stability landscape. In each of the three-dimensional graphs (upper panels), community compositions are shown on the two axes defined by a principal co-ordinate analysis (PCoA), while the vertical axis represented the inferred probability of observing specific community structure. Along the axis, a lower value (in blue) indicates a more probable community composition. In parallel, the statistical framework for inferring ecological dynamic regimes was applied to the dataset (lower panels). On each two-dimensional surface of PCoA, the time series of community dynamics are shown respectively for replicate communities (grey arrows) representing a dynamic regime. Representative trajectories, which summarize the main dynamic patterns of observed community assembly (Sánchez-Pinillos *et al*. 2023), are highlighted with yellow and green arrows (the green arrow is used as a reference to calculate dynamic dispersion). (A) Results for communities derived from a soil inoculum. (B) Results for communities derived from a freshwater inoculum.

## RESULTS

### Inoculum communities

The estimated number of prokaryotic cells contained in ×1 inocula were 2.59 × 10^6^ for the soil source community and 2.30 × 10^5^ for the freshwater source community (Table S3). The soil source community consisted of diverse taxa of bacteria such as *Pseudomonas*, *Stenotrophomonas*, *Burkholderia*, *Flavobacterium*, and *Klebsiella*, while the freshwater source community was dominated by *Pseudomonas* (Fig. S2). In total, 102 OTUs representing 50 genera and 62 OTUs representing 41 genera were observed in the soil and freshwater source communities, respectively (Fig. S3). The Shannon’s diversity of the OTUs were 3.53 for the soil source community and 2.03 for the freshwater source community.

### Community compositions of the experimental microbiomes

The experimental microbiomes deriving from the soil source community were constituted mainly by the five genera, *Klebsiella*, *Pseudomonas*, *Raoultella*, *Stenotrophomonas*, and *Serratia*, while those deriving from the freshwater source community were dominated by the three genera, *Pseudomonas*, *Yersinia*, and *Aeromonas* (Fig. 2B, Fig. S5). For both types of inoculum communities, the OTU-level compositions of experimental communities varied more conspicuously among replicate communities at higher dilution rates (e.g., at ×1/1000 dilution; Fig. 2). Such elevation of among-sample variation in community compositions with increasing dilution rate was observed as well at the ASV- and genus-level analyses (Fig. S4–5). In each combination of source microbiomes and inoculum dilution rates (i.e., inoculum setting), alpha diversity, as evaluated with the Shannon’s diversity or the number of OTUs, decreased through time in the microbiome experiment (Fig. S3).

### OTU-level differentiation among replicates

The analysis of the distributions of OTU abundance showed that multimodality increased with increasing level of stochasticity at the inoculation event (i.e., initial variation upon inoculation; Fig. 3D; Fig. S7). The trend was observed at all the time points for both the soil and freshwater inoculum experiments, while the level of multimodality was higher in the freshwater-inoculum experiment than in the soil-inoculum experiment (Fig. 3D).

The within-mode variation in the OTU abundance distributions significantly increased with increasing initial stochasticity (initial variation upon inoculation) at all the examined time points except for within-mode variation on Day 8 (Fig. 3E; Fig. S7A; see Table S6 for detailed results). In contrast to multimodality, the within-mode variation was consistently higher in the soil-inoculum experiment than in the freshwater-inoculum experiment (Fig. 3E). No significant correlation was observed between the two types of within-mode variation and multimodality (Fig. S7B).

### Community-scale differentiation among replicates

Multimodality in the distribution of dissimilarity among replicate communities (Fig. S8 and Table S8) was basically greater at higher dilution rates upon inoculation (Fig. 4D). Likewise, within-mode variation in the distribution of community dissimilarity was higher at higher dilution rates on both the soil and freshwater inoculum experiments (Fig. 4E).

In terms of temporal dynamics, the community-level multimodality values increased through time, although abrupt increase in multimodality was observed in the ×1 soil-inoculum setting on Day 4 (Fig. 4D). This exceptional pattern on Day 4 was attributed to the transient occurrences of replicate communities with abrupt structural changes: such peculiar community compositions disappeared on Day 6 (Fig. 2A).

In contrast to the overall pattern observed in the multimodality analysis, within-mode variation in community dissimilarity decreased over time in the freshwater-inoculum experiment (Fig. 4E). Within-mode variation in the soil-inoculum experiment exhibited no clear temporal trends (Fig. 4E).

### Ecological dynamic regimes

A series of the energy landscape analysis illustrated how stability landscape architecture could change depending on inoculum dilution rates (Fig. 5; Fig. S9-11). The community compositions of the same time points clustered on the inferred energy (stability) landscape under the ×1 inoculum condition (Fig. 5). In contrast, such clusters representing the same time points decayed with increasing inoculum dilution rates. At the highest dilution rate (×1/1000), data points are scattered on a complex topography of the stability landscape in both the soil- and freshwater-inoculum experiments (Fig. 5).

The ecological dynamic regime analysis further indicated that the directions of temporal community dynamics varied among replicate samples (dynamic regimes) more conspicuously at higher inoculum dilution rates (Fig. 5). In fact, dynamic dispersion (dDis) and dynamic beta diversity (dBD), which represented the extent to which the trajectories of community assembly varied among replicates, increased with increasing inoculum dilution rates (Fig. 6 left and middle; Fig. S12). As expected by the trends, dynamic evenness (dEve), which represented the continuity of variation within the trajectory space of community assembly, decreased with increasing inoculum dilution rates (Fig. 6 right).

**FIGURE 6.**
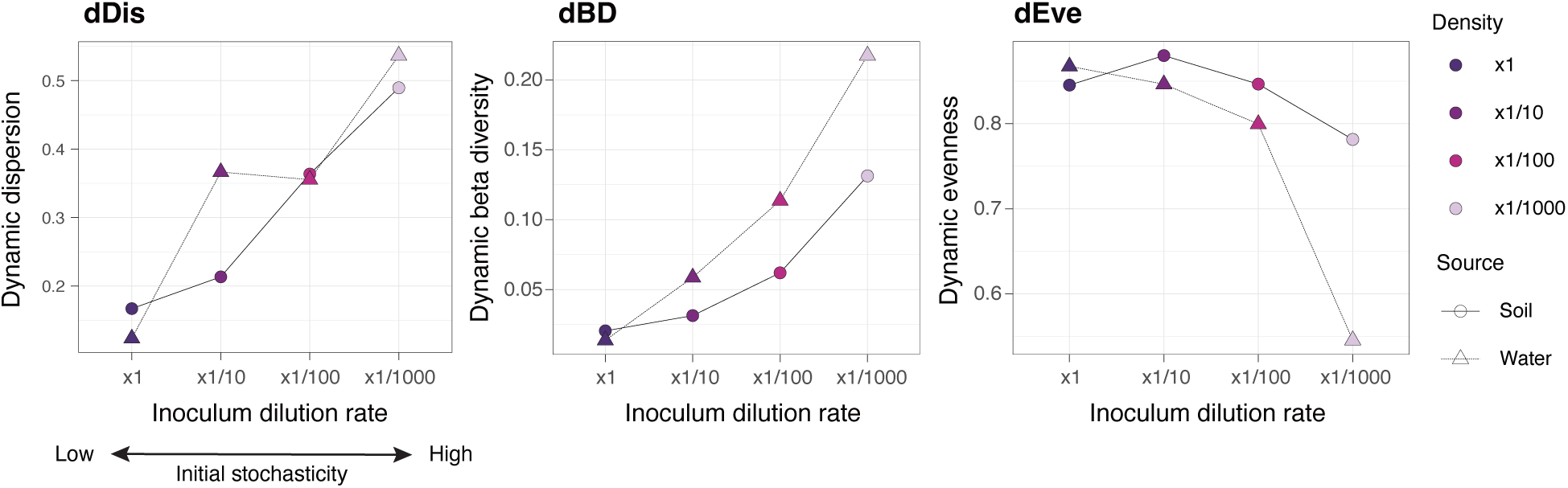
Divergence of ecological trajectories. Based on the ecological dynamic regime framework, community dynamics are characterized for each combination of source microbiome type (soil or freshwater) and inoculum dilution rate. A larger value of the dynamic dispersion (dDis) or dynamic beta diversity (dBD) metrics indicated greater differentiation in community-dynamics trajectories among experimental replicates. Meanwhile, a smaller dynamic evenness (dEve) value indicates greater divergence of ecological trajectories.

## DISCUSSION

By developing a highly parallelized system for tracking temporal community dynamics, we evaluated the roles of stochasticity at the foundation of communities. In the context of stochastic ecological processes, a number of studies have been designed to demonstrate how the order of species’ arrival may influence community assembly [i.e., priority effects (Zhou *et al*. 2013; Fukami 2015; Debray *et al*. 2022)]. Along with arrival orders, the size of arriving communities can critically impact early community assembly, eventually leading communities to alternative transient/stable states. With the multi-replicated experimental design (96 replicates/setting), we examined the extent to which slight stochastic differences could result in diverged community dynamics. Such quantitative evaluation of stochasticity and its ecological consequences provides a platform for deepening our understanding of community assembly through the lens of basic statistics and physics.

Our hypothesis on the relationship between initial stochasticity and the divergence of the subsequent community dynamics was supported by the microbiome experiment. The multi-replicated experimental design facilitated the quantitative evaluation of the extent to which the consequences of community dynamics were categorized into discrete community compositions. The multimodality analysis showed that the higher dilution rates (higher initial stochasticity) of founding communities resulted in greater levels of divergence into discrete OTU- and community-level patterns among replicate microbiomes (Figs. 2, 3D, and 4D). This result suggests that increased stochastic variation in community states at early community assembly facilitated splitting into alternative basins of attraction on a stability landscape (Fig. 1A right). In other words, the reproducibility of community dynamics *per se* is controllable if we can limit or augment stochasticity at early assembly by adjusting the size of arriving communities (Scheffer & Carpenter 2003; Siqueira *et al*. 2020; Lopes *et al*. 2024).

The quantitative approach of this study provided an opportunity for exploring the critical community size beyond which the reproducibility of community dynamics could be drastically changed. For both soil- and freshwater-derived experimental microbiomes, the differentiation of community compositions became striking when the total number of the founding prokaryotic cells was less than the order of 10^4^ (Fig. 4D). Meanwhile, differentiation into discrete community compositions was observable even at higher initial densities (lower inoculum dilution rates; Fig. 2). Such divergence in ecological outcomes may stem from slight differences among replicates in the balance between dominant species (Goldford *et al*. 2018). Alternatively, among-replicate variation in the presence or absence of rare species might have disproportionate influence on the divergence of ecological dynamics (Leitão *et al*. 2016; Jousset *et al*. 2017). In both scenarios, the strength of ecological drift would depend not only on the ratio of founding communities to their source communities (i.e., the dilution rates in this study), but also on the diversity and heterogeneity of the species pools (Santillan & Wuertz 2022; Le Moigne *et al*. 2023). In other words, when a species pool includes only a few species with identical abundance, highly consistent patterns of community assembly will be observed across replicate communities even with a low number of founding community size (e.g., 10^3^). As the level of stochasticity can be simulated based on species pool data (Fig. 3A), comparative analyses of source communities with varying alpha diversity will enable us to derive general quantitative insights into the relationship between founding community size and the reproducibility of community dynamics.

In addition to the multimodality index for inferring the presence of multiple basins on a stability landscape, we here implemented a statistical framework for quantifying the level of fluctuations within each basin of attraction. The approach integrating binomial distributions with normal distributions is broadly applicable when evaluating the extent to which ecological drift causes fluctuations in each species’ population dynamics (Fig. 3B). The analysis can be extended to the community level by combining multinomial distributions with normal distributions, estimating the level of fluctuations around the attractors of community compositions (Fig. 4A-B). By calculating the indices, we found that greater stochasticity at founding communities could result in stronger ecological drift within each basin (Figs. 3E and 4E). Meanwhile, an unexpectedly low level of “within-basin” variation was observed for the freshwater-derived experimental microbiomes with the highest dilution rate on Day 8 (Fig. 4E). Considering that the freshwater source community included lower alpha diversity than the soil source community (Fig. S3), the high initial stochasticity setting would have caused rapid divergence into alternative basins (Fig. 3D). Within each basin, stochastic processes (ecological drift) and deterministic processes such as competition between species with similar niches may have caused rapid reduction of species richness (Fig. S3), thereby limiting the range of variation across replicate communities representing the same basin (Chase & Myers 2011). The community structure of the ×1/1000 freshwater inoculum communities on Day 8 was characterized by the monodominance of alternative *Pseudomonas* OTUs (Fig. 2B), suggesting that competitive exclusion had occurred between *Pseudomonas* species. It remains an important challenge to explore the presence of any positive feedback mechanisms that could reinforce such monodominant states (e.g., antibiotics or pollutants inhibiting the growth of competitors) (Khare & Tavazoie 2015; Stubbendieck & Straight 2016).

In addition to the level of divergence among replicate communities, ecosystem functions may systematically change depending on initial community size. In the freshwater-inoculum experiment, the dominance of a single or a few *Pseudomonas* OTUs was observed at the highest dilution rate (×1/1000) as discussed above. In contrast, at the lowest dilution rate (×1), consistent patterns characterized by the coexistence of *Pseudomonas*, *Yersinia*, and *Raoultella* were observed across replicate communities (Fig. 2B). Since these genera differ in their functional profiles (Peng *et al*. 2024), the community-scale functioning of the three-genus coexistence state is expected to differ from the monodominance state of *Pseudomonas*. Thus, qualitative changes in ecosystem functions could result from quantitative changes in stochasticity during the initial stages of community assembly. In other words, the repertoires of alternative transient/stable states, which may differ in functional profiles, could drastically change depending on the stochasticity introduced at early assembly (Abbott & Nolting 2017).

This finding allows an alternative interpretation that the topography of stability landscapes, which is represented by the repertoires of basins, could change along the gradient of stochasticity (Framework 2). In this sense, the level of initial stochasticity *per se* could be interpreted as a hyperparameter determining assembly rules (Fig. 1B). Based on the alternative conceptual framework, we inferred how stability landscape architecture changed with increasing levels of initial stochasticity based on the energy landscape analysis (Suzuki *et al*. 2021). At the lowest level of initial stochasticity (×1 inoculum setting), community compositions at the same time points clustered together and shifted from hilltops to valleys on the inferred landscapes, suggesting that strong deterministic processes governed the community assembly (Fig. 5). In contrast, at the highest initial stochasticity level (×1/1000 inoculum setting), community compositions did not form clear clusters by time point, suggesting that elevated stochasticity drove divergent community dynamics among replicate samples. The divergence of the trajectories of community dynamics could be further overviewed in the framework of the ecological dynamic regime analysis (Sánchez-Pinillos *et al*. 2023) (Fig. 5). The statistical framework then allowed the quantification of the extent to which the trajectories of ecological dynamics varied across replicate communities. We then found that greater divergence of ecological trajectories could occur with stronger initial stochasticity (Fig. 6). These results illuminate the need to extend the concept of dynamically changing stability landscapes (Scheffer & Carpenter 2003; Suzuki *et al*. 2021) or ecological dynamic regimes (Sánchez-Pinillos *et al*. 2023, 2024) beyond discussions assuming fixed stability landscape architecture.

Although we developed a high-throughput way for systematically evaluating the ecological outcomes of stochasticity, the limitations of the present experimental design need to be acknowledged. First, our experiments with deep-well plates would have restricted the number of coexisting species in each replicate community. Within a small habitat, extinction of species is more likely to occur through ecological drift (Hubbell 2005). In addition, refugees of inferior competitors are less available in smaller habitats, promoting the dominance of superior species with similar ecological niches (Tilman 1994). Therefore, the experimental design should be extended to reveal how the relationship between initial stochasticity and diversity in ecological consequences can vary along the axis of culture volumes (Kram & Finkel 2014). Second, although we applied a mechanized experimental system, uncontrolled environmental factors worked as unexpected deterministic processes. For example, temperature might have fluctuated more severely in the outer than inner parts of a culture plate. Nonetheless, we were unable to detect such systematic spatial patterns of ecological consequences within each culture plate (Figs. S13-14). Third, the observation period may have been insufficient for observing the alternative stable states of community compositions. Even if the monodominant states observed on Day 8 in some inoculum settings (e.g., ×1/1000 freshwater) seemingly represent the destinations of community states, the resurgence of other bacteria enduring at very low densities may occur if sampling period is extended. It is generally difficult to distinguish alternative transient states from alternative stable states (Fukami & Nakajima 2011; Hayashi *et al*. 2024). Therefore, we did not use the term “alternative stable states” when interpreting the experimental results. In this sense, the terms “alternative trajectories” or “alternative dynamic regimes” can be broadly used without making controversial definitions of transience and steadiness in community dynamics (Scheffer & Carpenter 2003; Mayer & Rietkerk 2004; Angeler & Allen 2016). The ecological dynamic regime framework (Sánchez-Pinillos *et al*. 2023) provides a practical approach for quantitatively evaluating how community dynamics are differentiated based on initial stochasticity. Fourth, although we used fresh media in the experiment to examine the influence of stochasticity at the foundation of communities, such assembly in vacant environments rarely occurs in the wild. Consequently, it would be intriguing to redesign the experimental approach to evaluate how the size and arrival timing of immigrant communities can influence the divergence of resident communities’ dynamics.

Further insights into the interplay of deterministic and stochastic processes will not only enhance our basic understanding of community assembly but also advance applied sciences for controlling community-scale structure and functions. For example, in the use of plant-associated microbes in agriculture (Carlström *et al*. 2019), determining the founding community size of biostimulant microbiomes is essential for yielding stable functional benefits. Likewise, in the management of human gut microbiomes, reproducibility in the outcomes of microbiome transplantation (Green *et al*. 2020; Mingaila *et al*. 2023) would be quantitatively evaluated along the axis of introduced bacterial community size. In line with deepening the knowledge of stochastic processes, it is crucial to build a systematic understanding of deterministic processes when we develop frameworks for predicting and controlling community dynamics. If genomic information is available for the species constituting a community, the level of resource competition or metabolic exchange could be inferred at the community level (Zelezniak *et al*. 2015; Dukovski *et al*. 2021).

Thus, synthetic community experiments using genome-sequenced bacterial species or strains (Carlström *et al*. 2019; Vrancken *et al*. 2019), as well as shotgun metagenomic analyses of field-collected source microbiomes (Frioux *et al*. 2020; Fujita *et al*. 2023b), will help us evaluate the extent to which community-scale consequences can be predicted based on our understanding of deterministic processes. Systematic tests in microbial model systems will accelerate feedback between theoretical and empirical studies on temporal community dynamics, shedding new light on the assembly rules of macro-organisms.

## DATA AVAILABILITY

The 16S rRNA gene sequence data are available from the DNA Data Bank of Japan (DDBJ; accession number, Bioproject PRJDB35809) [to be released after acceptance of the paper]. The microbial community data and all the R scripts used for the statistical analyses are available at the GitHub repository (https://github.com/Ibuki-Hayashi/Founding-Community-Size-Governs-Divergent-Ecological-Trajectories) [to be released after acceptance of the paper].

## ACKNOWLEDGEMENTS

This work was financially supported by NEDO Moonshot Research and Development Program (JPNP18016), JST FOREST (JPMJFR2048), JST CREST (JPMJCR23N5), and CeLiSIS Program to HT. and JSPS Fellowship (24KJ1454) to IH.

## AUTHOR CONTRIBUTIONS

IH and HT designed the work. IH performed the experiments. IH analyzed the data with HT and MSP. IH and HT wrote the paper with MSP.

## COMPETING INTERESTS

HT is the founder and a director of Sunlit Seedlings Ltd. The other authors declare that the research was conducted in the absence of any commercial or financial relationships that could be construed as a potential conflict of interest.

