## Supporting information for "Stochastic Forces in Microbial Community Assembly: Founding Community Size Governs Divergent Ecological Trajectories"

**This PDF file includes:**

Supplementary methods
Supplementary Figures S1-14

**Supplementary Information included in a separate file:**

Supplementary Tables S1-7

**Supplementary Methods**

**DNA extraction**

To extract DNA from each culture sample, 5 μL of the collected aliquot was mixed with 1 μL lysozyme solution [50 mg/ml lysozyme (Sigma), 20 mM Tris-HCl (pH 8.0), 2 mM EDTA] and the mixed solution was incubated at 37 ºC for 2 hours. After adding proteinase K solution [1/30 (v/v) Proteinase K (Takara), 20 mM Tris-HCl (pH 8.0), 2 mM EDTA], the aliquot was incubated at 55 ºC for 3 hours and 95 ºC for 10 min. The solution was then vortexed for 10 minutes to increase DNA yield.

**PCR and DNA sequencing**

For the samples of the experimental microbiomes, prokaryote 16S rRNA V4 region was PCR-amplified with the forward primer 515f fused with 3–6-mer Ns for improved Illumina sequencing quality (Lundberg *et al.* 2013) and the forward Illumina sequencing primer (5’- TCG TCG GCA GCG TCA GAT GTG TAT AAG AGA CAG- [3–6-mer Ns] – [515f] -3’) and the reverse primer 806rB fused with 3–6-mer Ns and the reverse sequencing primer (5’- GTC TCG TGG GCT CGG AGA TGT GTA TAA GAG ACA G [3–6-mer Ns] - [806rB] -3’) (0.2 μM each). The buffer and polymerase system of KOD One (Toyobo) was used with the temperature profile of 35 cycles at 98 ºC for 10 s, 55 ºC for 5 s, 68 ºC for 1 s. To prevent generation of chimeric sequences, the ramp rate through the thermal cycles was set to 1 ºC/sec (Stevens *et al.* 2013). Illumina sequencing adaptors were then added to respective samples in the supplemental PCR using the forward fusion primers consisting of the P5 Illumina adaptor, 8-mer indexes for sample identification (Hamady *et al.* 2008) and a partial sequence of the sequencing primer (5’- AAT GAT ACG GCG ACC ACC GAG ATC TAC AC - [8-mer index] - TCG TCG GCA GCG TC -3’) and the reverse fusion primers consisting of the P7 adaptor, 8-mer indexes, and a partial sequence of the sequencing primer (5’- CAA GCA GAA GAC GGC ATA CGA GAT - [8-mer index] - GTC TCG TGG GCT CGG -3’). KOD One was used with a temperature profile of 8 cycles at 98 ºC for 10 s, 55 ºC for 5 s, 68 ºC for 5 s (ramp rate = 1 ºC/s). The PCR amplicons of the samples were then pooled after a purification/equalization process with the AMPureXP Kit (Beckman Coulter). Primer dimers, which were shorter than 200 bp, were removed from the pooled library by supplemental purification with AMPureXP: the ratio of AMPureXP reagent to the pooled library was set to 1 (v/v) in this process. This library was further purified with E-gel SizeSelect 2 (Invitrogen) and then ca. 440-bp DNA fragments was selectively obtained. The sequencing libraries were processed in an Illumina MiSeq sequencer [271 forward (R1) and 31 reverse (R4) cycles; 20% PhiX spike-in].

For the source microbiome sample, the prokaryote 16S rRNA V4 region was also amplified. To estimate concentrations of 16S rRNA genes included in the inoculum, a quantitative amplicon sequencing platform was applied by introducing five “standard DNA” fragments with controlled concentrations to the PCR master mix solution of the first PCR process as detailed elsewhere (Ushio *et al.* 2018). The standard DNAs were used for the *in-silico* calibration of 16S rRNA gene concentrations in the target sample after sequencing as detailed in the previous study (Fujita *et al.* 2023).

**Bioinformatics**

In total, 66,302,381 sequencing reads were obtained with the Illumina sequencing. The raw sequencing data obtained in the Illumina sequencing were converted into FASTQ files using the program bcl2fastq 1.8.4 distributed by Illumina. The output FASTQ files were then demultiplexed with the program Claident v0.9. 2022.01.26. The sequencing reads were subsequently processed with the program DADA2 (Callahan *et al.* 2016) v.1.18.0 of R 4.2.2 to remove low-quality data. The molecular identification of the obtained amplicon sequence variants (ASVs) was performed based on the naive Bayesian classifier method (Wang *et al.* 2007) with the SILVA v.138.1 database (Quast *et al.* 2013).

**Overview of the community data**

The rarefaction curves indicating relationship between the number of sequencing reads and the number of ASVs were drawn using the vegan 2.6.11 package (Oksanen *et al.* 2025) of R 4.4.0. In the sequencing of source (inoculum) microbiome samples, the diversity of microbial ASVs reached plateaus along the axis of the number of sequencing reads (Fig. S1). The data of sequencing read counts were converted to those of DNA concentrations based on the calibration procedure detailed in the “Bioinformatics” and “Estimating initial variation upon inoculation” subsections of the main manuscript.

In the sequencing of experimental culture samples, the diversity of microbial ASVs also reached plateaus along the axis of sequencing read counts (Fig. S1). Given the rarefaction curves, the dataset of experimental culture samples was rarefied to 3,000 reads per sample with the "rrarefy" function of the R vegan package. Of the 3,072 samples (8 inoculum conditions × 96 replicates × 4 time points), 3,063 samples with more than 3,000 reads were used in the following pipeline. After screening for the replicate communities for which sequencing data were available for all the four time points, 3,040 samples were subjected to the following statistical analyses. In total, 490 prokaryote ASVs belonging to two kingdoms, 26 classes, 64 orders, 88 families, and 147 genera were detected. Then, the ASVs were re-clustered into operational taxonomic units (OTUs) with a threshold similarity of 99% using the program VSEARCH v2.15.2 (Rognes *et al.* 2016), yielding, in total, 337 OTUs.

For a simple evaluation of the temporal fluctuation of the community, *α*-diversity and temporal variability of each community were quantified. Two types of *α*-diversity indices, OTU richness and Shannon-Wiener diversity, were calculated for each sample. Then, two types of indices, Jaccard dissimilarity and Bray-Curtis dissimilarity between adjacent days, were used to evaluate temporal variability. All metrics were calculated with the vegan package.

**Energy landscape analysis**

To infer the stability landscape architecture of the experimental microbiomes, we applied the statistical framework of energy landscape analysis, which captures complex systems' behavior based on the Ising model of statistical physics (Suzuki *et al.* 2021; Masuda *et al.* 2025; Toju et al., *in revision*). The statistical framework has been applied to neuroscience (Watanabe *et al.* 2014) and ecology (Suzuki *et al.* 2021; Fujita *et al.* 2023; Kadoya *et al.* 2025), giving insights into the multi-stable states of systems. When applied to ecological community data (tutorials and R codes of energy landscape analyses are available at <https://github.com/kecosz/rELA>), the probability of observing a specific community state [$P\left( \vec{\sigma}^{\left( k \right)} \right)$] is expressed as:

$P\left( \vec{\sigma}^{\left( k \right)} \right)={e^{-E\left( \vec{\sigma}^{\left( k \right)} \right)}}/Z$,

$Z=\sum_{i=1}^{2^{S}} e^{-E\left( \vec{\sigma}^{\left( k \right)} \right)}$,

where $\vec{\sigma}^{\left( k \right)}=\left( {\sigma_{1}}^{\left( k \right)}, {\sigma_{2}}^{\left( k \right)},\ldots,{\sigma_{S}}^{\left( k \right)} \right)$ is a community state vector of *k*-th sample and $S$ is the total number of the taxa examined (e.g., the number of OTUs, species, genera, or families in the input data). Within the community state vector, ${\sigma_{i}}^{\left( k \right)}$ is a binary variable that indicates presence (1) or absence (0) of taxon *i*: i.e., there are a total of $2^{S}$ community states. When input community matrix is defined, the $E\left( \vec{\sigma}^{\left( k \right)} \right)$ part of the equation is fitted based on an extended pairwise maximum entropy model defined as follows:

$E\left( \vec{\sigma}^{\left( k \right)} \right)=-\sum_{i=1}^{S} h_{i} {\vec{\sigma}^{\left( k \right)}}_{i}-\sum_{i=1}^{S} \sum_{j=1, i\neq j}^{S} {J_{ij} {\vec{\sigma}^{\left( k \right)}}_{i} {\vec{\sigma}^{\left( k \right)}}_{j}}/2$,

where $h_{i}$ represents the net effect of implicit abiotic factors, by which *i*-th taxon is more likely to present ($h_{i}>0$) or not ($h_{i}<0$), and $J_{ij}$ represents the pattern of co-occurrence between *i*-th and *j*-th taxa. Since the logarithm of the probability of a community state is inversely proportional to $E\left( \vec{\sigma}^{\left( k \right)} \right)$, a community state having lower *E* is observed more frequently. Note that the "energy" metric (*E*) does not correspond in any way to the physical form of energy: it is specifically defined with the above equation in energy landscape analysis (Suzuki *et al.* 2021). Based on the statistical model, community states that show lower $E\left( \vec{\sigma}^{\left( k \right)} \right)$ values than all adjacent community states within an assembly graph are explored, inferred as attractors of community dynamics (Suzuki *et al.* 2021).

In applying energy landscape analysis to our dataset, the rELA 0.70 library (<https://github.com/kecosz/rELA>) of R was used. The original community data were converted into binary input data using the following read count threshold: OTUs that accounted for at least $4.32 \times{10}^{-4}$% of the total reads ($=5$ reads $/$ $(384$ samples $\times3000$ reads$)$ ) and appeared in at least 5 out of 384 samples started with the same inoculum condition were used as input data.

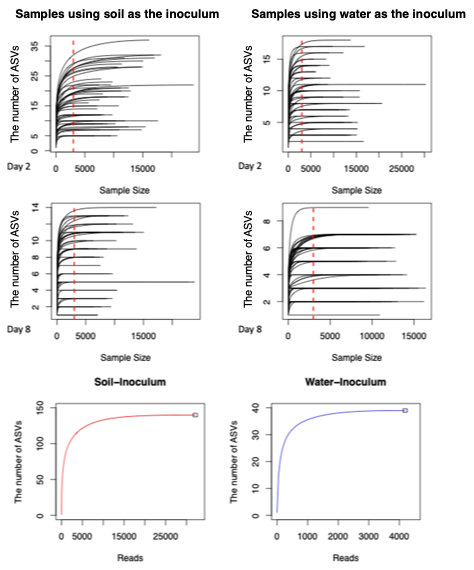

**B**

**A**

**Figure S1 |** Rarefaction curve of culture samples and inoculum samples. Relationships between the number of sequencing reads and the number of detected prokaryote ASVs are shown. (A) Rarefaction curves of 50 samples randomly selected from the pool of the samples with 3,000 or more sequencing reads are shown in each panel. The data of Day 2 and Day 8 are shown. (B) Rarefaction curves of the soil and freshwater source (inoculum) microbiomes.

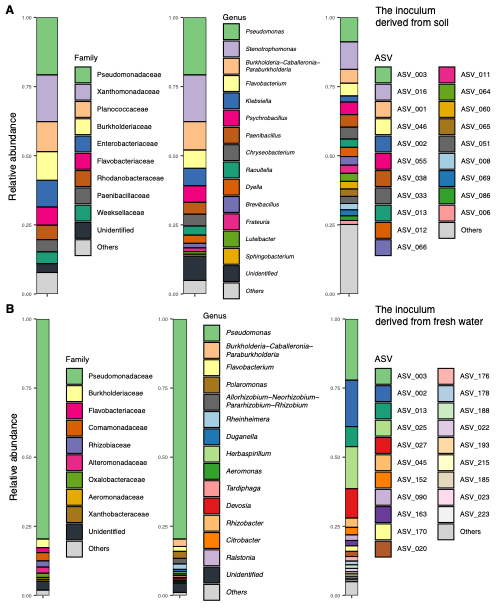

**Figure S2 |** Taxonomic compositions of soil and freshwater source (inoculum) microbiomes. For each of the soil (A) and freshwater (B) source microbiomes, community compositions are respectively shown at the family, genus, and ASV levels.

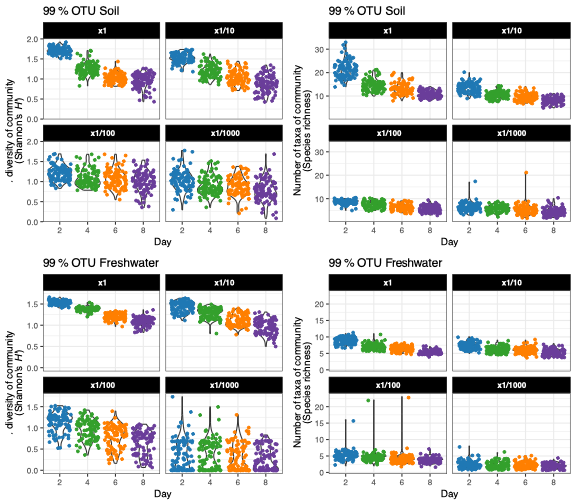

**Figure S3 |** Alpha diversity of the experimental microbiomes. Temporal changes in the Shannon’s diversity index (left) and richness (right) of 99% OTUs are shown for the soil (top) and freshwater (bottom) experimental microbiomes. The results of Student’s *t*-tests comparing each pair of time points are presented in Table S3.

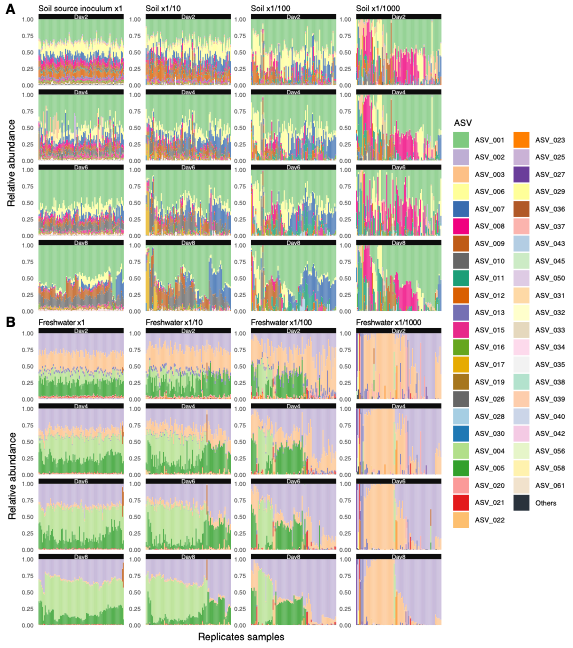

**Figure S4 |** Overview of among-replicate variation in community structure (ASV-level results). (A) Experiment with soil inoculum microbiome. Temporal changes in the ASV-level community compositions (relative abundance) are shown. For each inoculum dilution rate, replicate communities were ordered based on the results of unweighted pair group method with arithmetic mean (UPGMA) analyses performed on Day 8. (B) Experiment with freshwater inoculum microbiome.

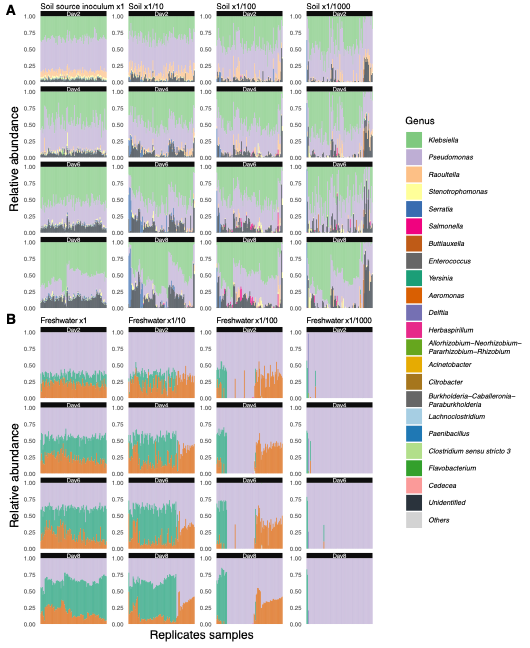

**Figure S5 |** Overview of among-replicate variation in community structure (genus-level results). (A) Experiment with soil inoculum microbiome. Temporal changes in the genus-level community compositions (relative abundance) are shown. For each inoculum dilution rate, replicate communities were ordered based on the results of unweighted pair group method with arithmetic mean (UPGMA) analyses performed on Day 8. (B) Experiment with freshwater inoculum microbiome.

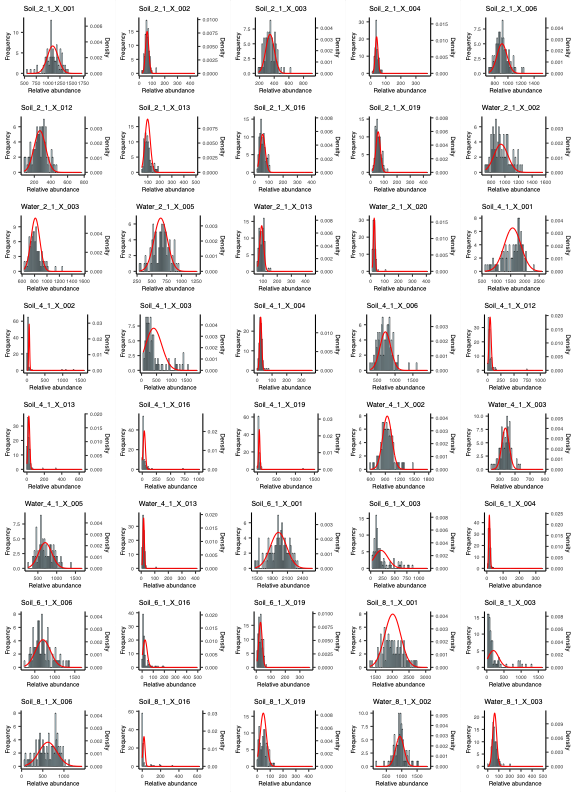

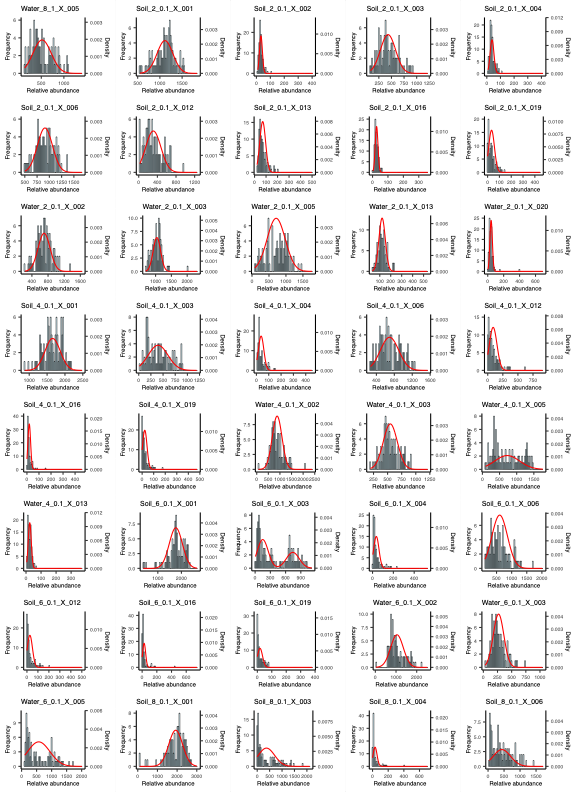

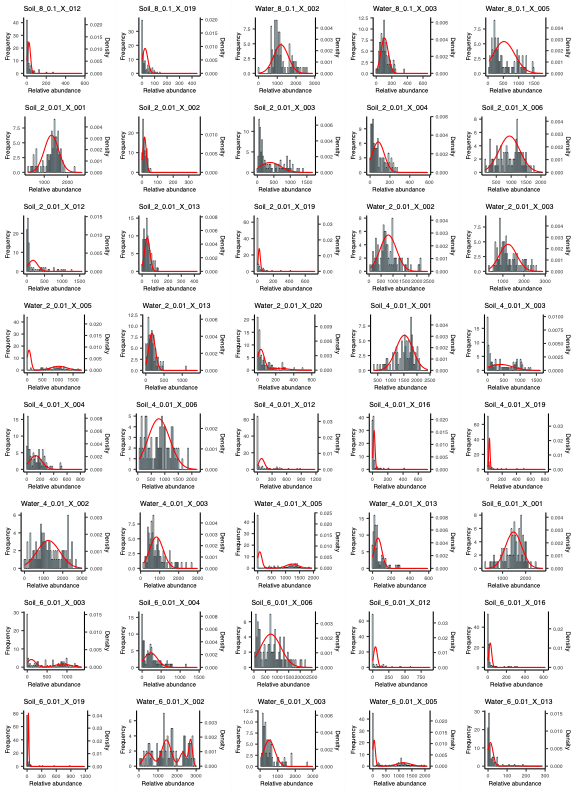

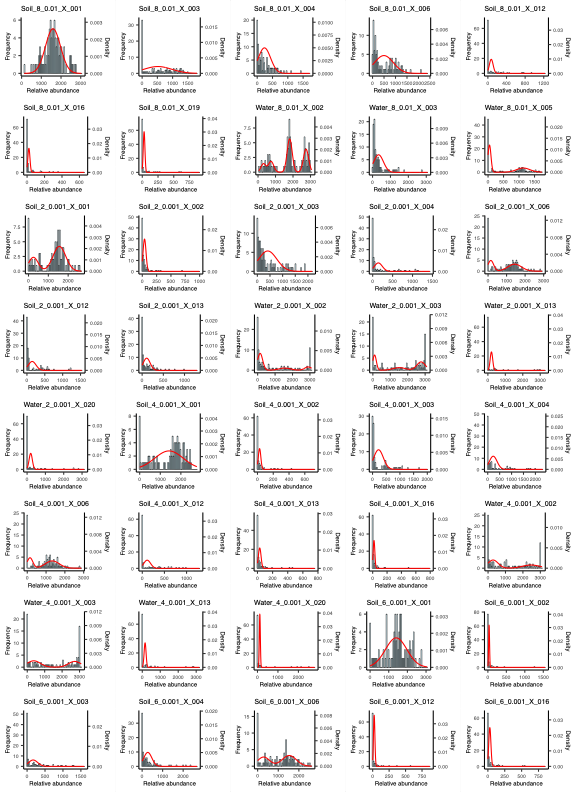

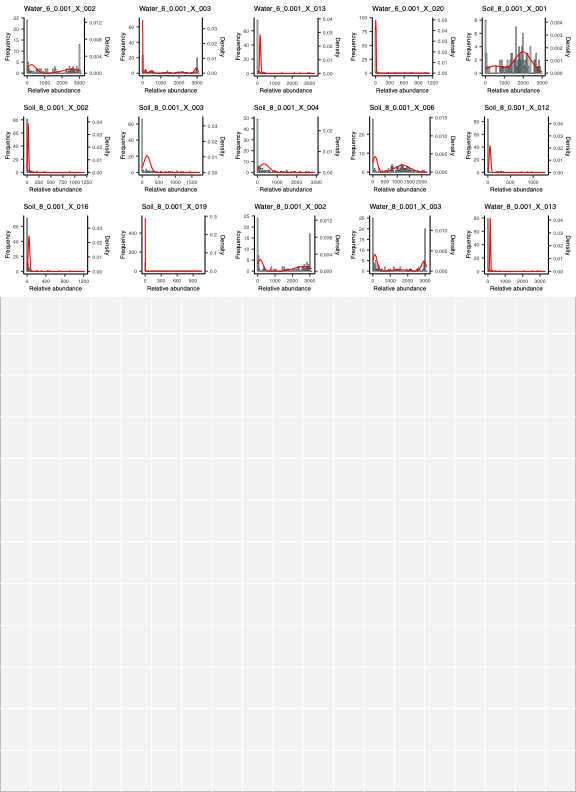

**Figure S6 |** Histograms of OTU abundance across replicate communities. Each histogram is overlaid with predictions from the mixture model (see Materials & Methods). The red curve in each panel represents the model’s prediction. Each panel shows the abundance distribution for a specific combination of inoculum source, sampling date, dilution rate, and OTU. The combination is indicated at the top of each panel (e.g., in “Soil_2_1_X_001”, the data represent a non-diluted soil-derived inoculum sampled on Day 2, with a focus on OTU X_001).

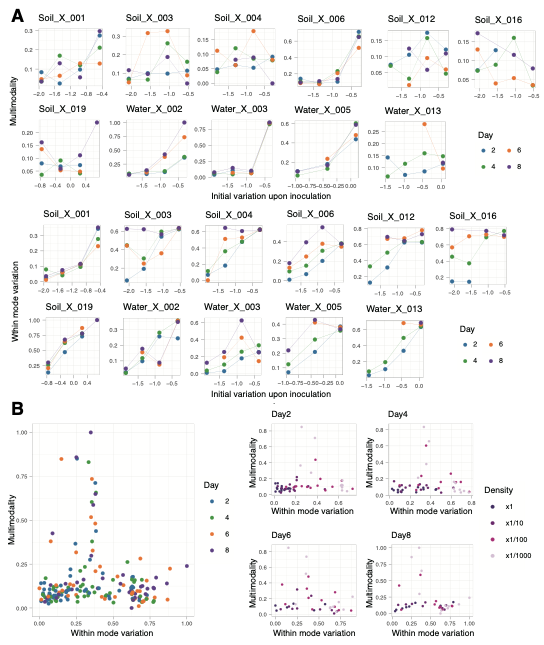

**Figure S7 |** Supplementary results on the analysis of multimodality and within-mode variation. (A) Temporal changes of multimodality and within-mode variation (OTU-level). The temporal changes in the relationships between the relationship between the multimodality of each taxon and the initial variation upon inoculation (upper) and the within-mode variation of each 99% OTU and the initial variation upon inoculation (lower) are shown in each panel. Only OTUs observed under at least three different initial density conditions within at least one day are shown. The used type of inoculum resource and focused OTU is indicated at the top of each panel (e.g., in “Soil_X_001”, the data represent a soil-derived inoculum, with a focus on OTU X_001). (B) The relationship between the within-mode variation and the multimodality. The relationship between the within-mode variation and the multimodality of each 99 % OTU is shown, with colors indicating different dates. Subplots at the left display the main plots separated by date, where colors represent different dilution rate.

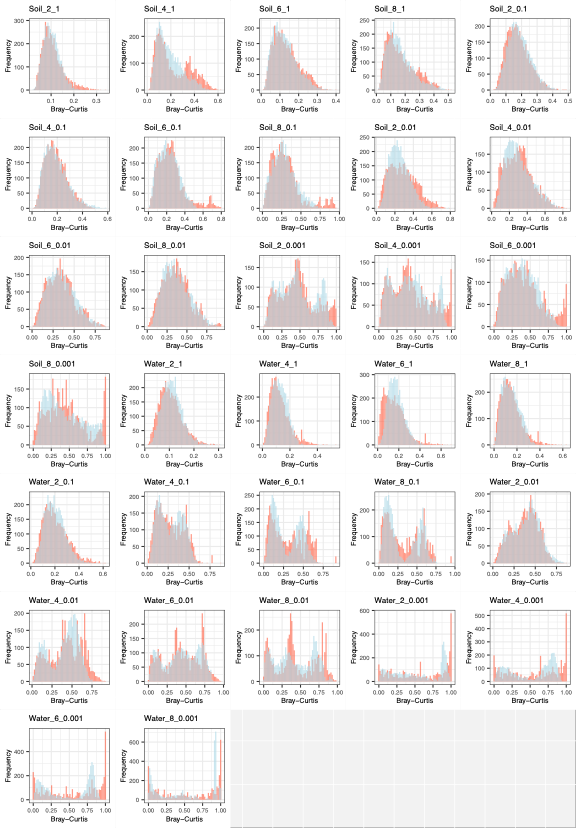

**Figure S8 |** Community dissimilarity histograms. Community dissimilarity histograms overlaid with predictions from the mixture model (see Materials & Methods) are shown (Theses panels are the full results of Fig. 4C). In each panel, the red histogram represents the actual data, while the blue histogram shows the distribution calculated from the mixture model. The overlap between actual and simulated distributions are indicated in grey. Each panel displays the community dissimilarity distribution for a specific combination of inoculum source, sampling date, and dilution rate. The combination is indicated at the top of each panel (e.g., in “Soil_2_1”, the data represent a non-diluted soil-derived inoculum sampled on Day 2).

**Figure S9 |** Detailed presentation of the inferred topographies of stability landscapes. For better visualization, the energy landscapes in Figure 5 are shown in enlarged form. The panels are organized horizontally by inoculum source and vertically by the dilution rate of the inoculum microbiome.

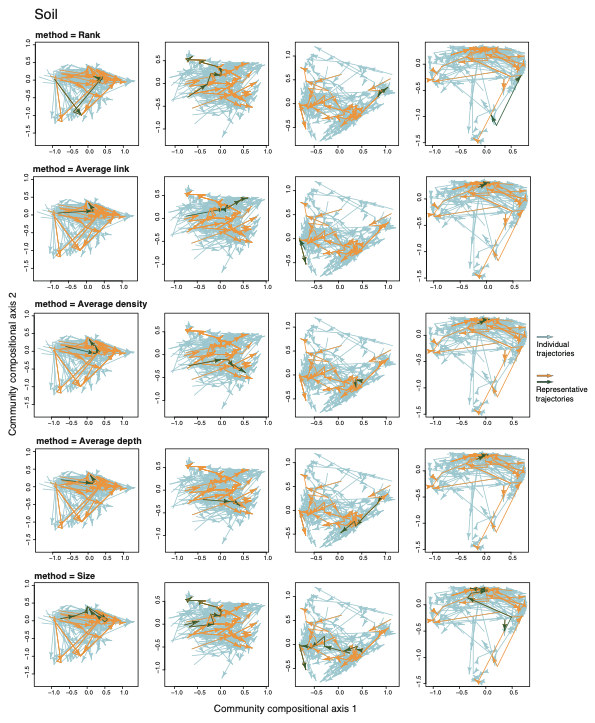

**Figure S10 |** Criteria for selecting reference trajectories in the analysis of ecological dynamic regimes (soil-derived microbiomes). The reference trajectories (green arrows) selected with each of the five criteria are shown in the PCoA plot of soil-derived communities analyzed using the framework of ecological dynamic regimes in Figure 5. The result of the method using the average depth is also shown in Figure 5. The method used to select the reference is indicated at the top of each panel. Panels are arranged horizontally by dilution rate.

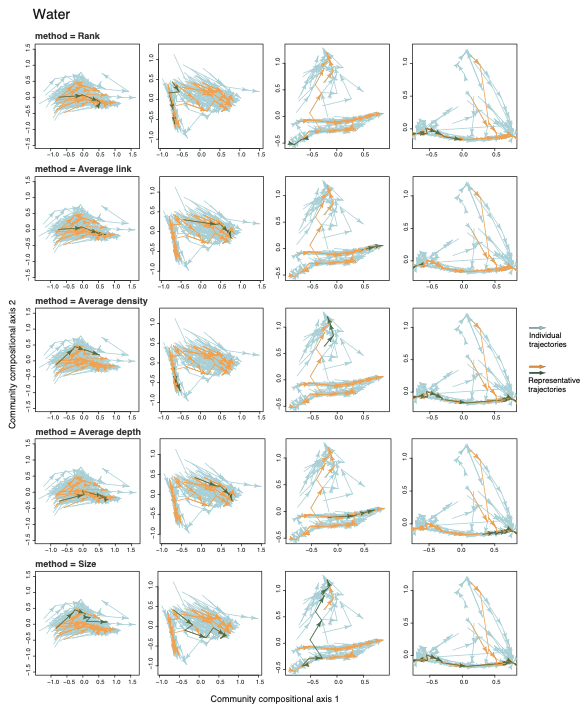

**Figure S11 |** Criteria for selecting reference trajectories in the analysis of ecological dynamic regimes (freshwater-derived microbiomes). The reference trajectories (green arrows) selected with each of the five criteria are shown in the PCoA plot of freshwater-derived communities analyzed using the framework of ecological dynamic regimes in Figure 5. The result of the method using the average depth is also shown in Figure 5. The method used to select the reference is indicated at the top of each panel. Panels are arranged horizontally by dilution rate.

**Figure S12 |** Dynamic dispersion metrics based on alternative criteria. In each panel, a reference trajectory selected by each method was used in the calculation of the dynamic dispersion. Except when using the size as the selection criterion, consistent patterns of increasing dynamic dispersion values with increasing dilution rates was observed.

**Figure S13 |** Spatial distributions of community structure within the culture plate (soil-derived communities). To assess whether the spatial arrangement of wells within the plate affects community structure, we conducted a clustering analysis of replicate communities. For each dilution rate of the soil-derived microbiome, we applied *k*-means clustering followed by silhouette coefficient analysis to determine the optimal number of clusters. After selecting the number of clusters *k*, we used principal coordinate analysis (PCoA) to reduce the dimensionality of community composition data and visualized the results, with colors representing different clusters. The spatial positions of replicate communities assigned to each cluster are shown on the right panels indicating the spatial organization within the culture plate.

**Figure S14 |** Spatial distributions of community structure within the culture plate (freshwater-derived communities). To assess whether the spatial arrangement of wells within the plate affects community structure, we conducted a clustering analysis of replicate communities. For each dilution rate of the freshwater-derived microbiome, we applied *k*-means clustering followed by silhouette coefficient analysis to determine the optimal number of clusters. After selecting the number of clusters *k*, we used principal coordinate analysis (PCoA) to reduce the dimensionality of community composition data and visualized the results, with colors representing different clusters. The spatial positions of replicate communities assigned to each cluster are shown on the right panels indicating the spatial organization within the culture plate.
